# Seasonal changes in the population structure of *Australpavlovskyella gurneyi*, the kangaroo soft tick, associated with seasonal changes in the wallowing behaviour of the *Osphranter rufus*, the red kangaroo, and the weather

**DOI:** 10.64898/2026.01.21.700930

**Authors:** Bernard M. Doube, Stephen C. Barker

## Abstract

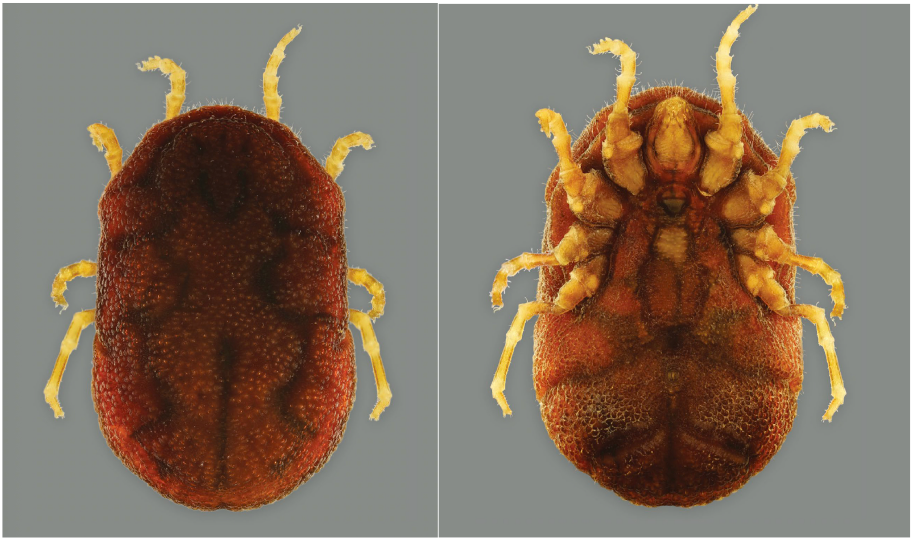

The kangaroo soft tick, *Australpavlovskyella gurneyi* (Warburton, 1926), is found in sandy depressions (‘wallows’), under desert shade trees, formed by the activity of the red kangaroo, *Osphranter rufus*, resting under shade trees (https://youtu.be/AYLoqqPsifc). The field biology of the tick was examined on Moralana Station in arid mid-north, South Australia, between February 1969 and March 1971. The age of kangaroo dung in wallows showed that kangaroos visited wallows regularly during the hot summer and infrequently during the cooler months. All nymphal instars and adults were present at all times of the year in kangaroo wallows, but only a small proportion of the ticks present was trapped on any one occasion. Ticks were abundant in large kangaroo wallows under trees with dense shade, but scarce under smaller trees with sparse shade. The short-lived larvae were present only during spring and early summer, indicating that the long-lived female ticks bred only during spring and early summer. Laboratory tests showed that field-collected adult female ticks entered reproductive diapause from January to August (mid-summer to late-winter). Ticks placed in kangaroo wallows survived for at least one year without food. On Moralana Station, the population of first-instar nymphs increased in summer and subsequently the population of second-instar nymphs increased in early autumn, indicating that a life cycle could be completed in 2–3 years.

**Highlights:**

- The seasonal biology of *Australpavlovskyella gurneyi*, found in sandy depressions ‘wallows’ formed by the activity of the red kangaroo, under sparse semi-arid desert shade trees was examined for the first time.
- Engorged ticks placed in kangaroo wallows survived for at least one year without food.
- In this environment, the entire life cycle could be completed in 2–3 years.

## 1. Introduction

Fourteen species of Argasidae are known in Australia (Barker et al., 2014). One of these, the kangaroo soft tick, *Australpavlovskyella gurneyi*, is renowned for its ability to live in the extremely harsh deserts of Central Australia (Henry, 1938; Browning, 1962; Roberts, 1970; Doube, 1972; Barker and Barker, 2023) where its principal host is the red kangaroo, *Osphranter rufus*. The tick is found in sandy depressions (‘wallows’) under desert shade trees that have been formed by the activity of the red kangaroo as it rests in the shade during hot summer days. Ticks burrow into the sand to avoid desiccation (refer to video below). Kangaroo wallows are relatively discrete and the number of wallows under each shade tree usually increases as the area shaded increases; with individual wallows separated by an area of undisturbed soil or by shrubby vegetation. Tick-infested wallows are found only in regions with relatively sparse tree cover (Browning, 1962). Due to the movements of the kangaroos, the soil in wallows is usually soft and friable and so *A. gurneyi* can burrow into the soil with ease.

–YouTube link to movie of ticks burrowing into sand – (https://youtu.be/AYLoqqPsifc).

Recent publications concerning ticks of the subfamily Ornithodorinae have been concerned primarily with their role in tick-induced toxicosis (Mans et al., 2002b); in the epidemiology (Reck et al., 2013) and transmission of viruses such as African swine fever virus (Oleaga et al., 1990), *Rickettsia* (Numan et al., 2022), Alkhuma hemoragic fever (Charrel et al., 2007) and *Borrelia* relapsing fever (Souidi et al., 2014); and their systematics (Boinas et al., 2014; Mans et al., 2019, 2021, 2025; Muñoz-Leal et al., 2020; Vázquez-Guerrero, 2023; Barker et al., 2025). *Ornithodoros* (*Pavlovskyella*) *gurneyi* (Warburton, 1926) is so different to other *Ornithodoros* species that a new genus, *Australpavlovskyella* Barker, 2025, was created for it recently (Barker et al., 2025).

The ecology of Ornithodorinae ticks in the field has received sparse attention apart from Neville, (1964), Adeyeye and Butler (1989), Doube (1972), Dupraz et al. (2017) and Vial et al. (2018). Neville (1964) studied the ecology of *O. savignyi* in South Africa. Mans coauthored extensive publications on the genus *Ornithodoros*, especially concerning the toxicity of *O. savignyi* and, more generally, on the role of soft ticks as parasites and vectors (e.g. Mans et al., 2022). Vial et al. (2018) developed spatial analyses of suitable habitats of *Ornithodoros* ticks in the Palearctic region. They suggested key survival criteria based upon spring and summer temperatures and seasonal rainfall patterns but excluded host availability and the effects of suitable habitat. Dupraz et al. (2017) demonstrated a temporal dynamic, with the simultaneous appearance of juveniles of the yellow-legged gull (*Larus michahellis*) and juveniles of the argasid *Alectorobius maritimus* but there were no among-nest differences in population structure. Adeyeye and Butler (1989, 1991) analysed the population structure and seasonal intra-burrow movement of *Ornithodoros turicata* in gopher tortoise burrows using CO_2_ as a lure and showed low inter-burrow movement but specific depth preferences within the burrows. Here we analyse the seasonal population biology of an Australian argasid tick in relation to temperature, rainfall, habitat preferences and host behaviour.

There are two strains of the kangaroo soft tick, a plains strain which feeds on the red kangaroo, *Osphranter rufus*, and the western grey kangaroo, *Osphranter fuliginosus*, and inhabits wallows in sparsely wooded open country, and a cave strain which inhabits caves in hilly country and feeds on the common wallaroo, *Osphranter robustus*, and the yellow-footed rock wallaby, *Petrogale xanthopus*, when they rest in caves during the heat of a summer day (Doube, 1975a). The present study concerns only the plains strain.

The life stages of *A. gurneyi* comprise a 6-legged larvae, 8-legged nymphs (between 3 and 5 instars), and adults (Doube, 1975b), which are very long lived (> several years) (Doube, 1972). Adult females display a reproductive diapause which is induced by short photoperiod and high temperature (Doube, 1975b, 1975c). Adults and nymphs can rehydrate from unsaturated air, thereby increasing longevity in the absence of a suitable host (Doube, 1972). The engorged larvae and nymphs display a circadian rhythm of detachment in the laboratory on rats: 2–12 days after attachment, the larvae and nymphs detached from their rat-hosts during the middle of the photophase (daylight). This would apparently optimise the chances that the engorged larvae and nymphs would drop off into a wallow occupied by a host kangaroo, thereby achieving both dispersal and an improved probability of finding another host (Doube, 1975d).

The red kangaroo rests by day and feeds by night, but whether it rests in a shaded wallow or in an unshaded area depends on the temperature of the day. On cold days kangaroos bask in the sun in areas sheltered from wind (e.g. behind clumps of bushes) whereas on hot days they shelter from the sun under shady trees (Frith and Calaby, 1969). Since this behaviour is a thermo-regulatory mechanism, the frequency of use of any tree species depends on the quality of shade provided and the temperature of the day.

In the present paper, we show that the plains race of the kangaroo soft tick, *A. gurneyi*, lives in dry Australian desert regions in the sandy soil of kangaroo wallows under sparsely distributed shade trees that were periodically occupied by the red kangaroo during hot summer days. All instars of the tick fed on the kangaroo, but the females bred largely during spring to early summer and, in the study area, the egg-to-adult life cycle apparently took at least 2–3 years. Seasonal changes in the structure of the tick population reflected the seasonal activity patterns of the host kangaroo.

## 2. Materials and Methods

### 2.1. Study area

The study area, Moralana Station (latitude –31.52759, longitude 138.2775), is located about 360 km north of Adelaide, SA, between the Flinders Ranges and Lake Torrens (Fig. 1). Both *A. gurneyi* and red kangaroos were plentiful, and a section between two sand dunes was selected as the main location for the field study. The landscape comprised a series of east–west sand dunes (populated by solitary bullock bushes (*Alectryon* spp.) or black oaks (*Casuarina* spp.). The slopes between the dunes were populated with solitary mulga trees (*Acacia aeneura*) and a small bushy *Acacia* sp. (plains acacia), and prickly acacia (*Acacia victoriae*) occurred on the mud pans between the dunes. Bullock bushes provided the densest shade whereas black oaks provided the most variable shade quality. The study was between February 1969 and March 1971.

**Fig. 1.**
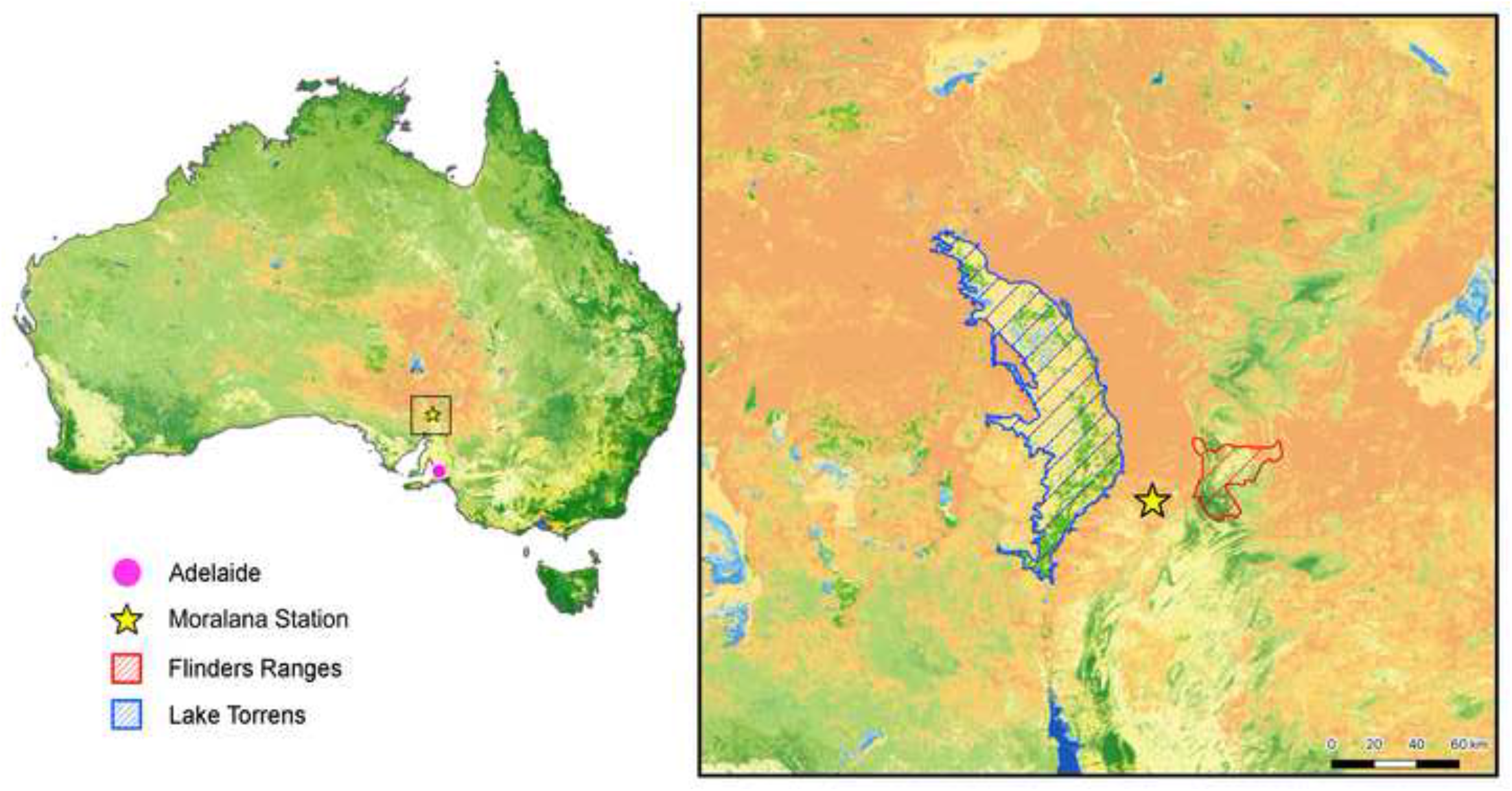
Location of the study area.

### 2.2. Sampling tick populations

*Australpavlovskyella gurneyi* was collected from Moralana Station field by using CO_2_ derived from dry ice as a lure (rather than by sifting sand) as for other tick species (Neville, 1964). Other methods (warm tanned kangaroo skin, vibrating the soil with a mallet) failed to lure ticks to the soil surface. Each CO_2_ trap consisted of a petri dish buried in the sandy wallow such that the lip of the petri dish was level with the surface of the sand. A lump of dry ice was placed on an elevated platform in the centre of each petri dish. Ticks emerged from the sand within minutes of the CO_2_ lure being placed and walked across the sand and fell into the petri dish, from which they could not escape. Ticks continued to arrive in the trap for as long as the dry ice lasted, generally some hours. One day of tick-trapping, did not remove all of the ticks that were in a wallow. This was confirmed in laboratory observations in which newly moulted ticks were placed in an artificial wallow and the ticks lured to the surface on the following day. About 30% of all stages failed to respond to a CO_2_ lure. Similar behaviour was observed in the sand tampan, *Ornithodoros savignyi*, in South Africa (Neville, 1964).

The standard trapping-protocol comprised a pair of pitfall traps in each wallow which were baited in the morning and cleared in the afternoon. This approach was based on an initial study in which traps were set in pairs of kangaroo wallows under a series of trees. The total number of ticks collected per tree ranged from 35 to 659 but the age structure (relative numbers of each life stage (ie instar)) was very similar in each of the paired traps whereas the age structure varied widely among trees. On this basis it was decided to sample under a moderate number of trees and to set two traps per wallow on each sampling occasion.

The ticks from each trap were placed in a container and taken to the laboratory, where the length of all nymphs and adults was measured and the number of larvae counted. A frequency distribution of the lengths of the ticks allowed recognition of first-, second- and third–fifth-instar nymphs, providing six categories (larvae, 1NN, 2NN, 3^-^5NN and adult males and females) (Fig. 2).

**Fig. 2.**
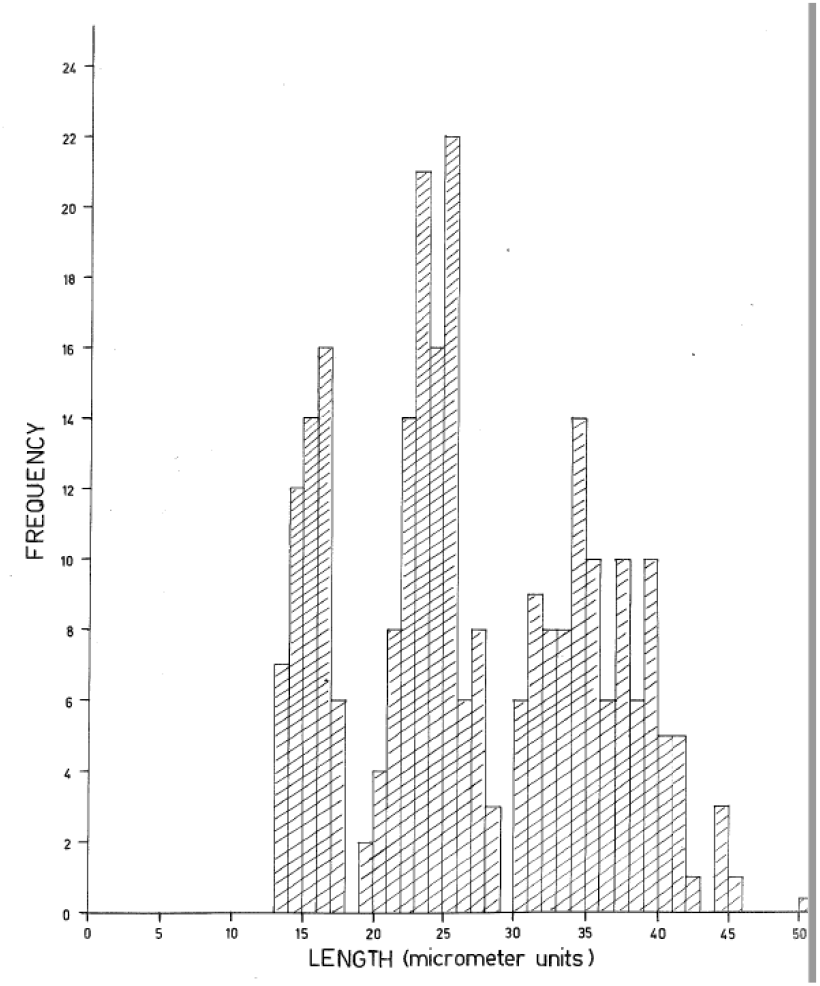
Frequency distribution of lengths of nymphs of *Australpavlovskyella gurneyi* (first-instar to fifth-instar).

### 2.3. Age structure of tick populations

The number of large ticks (adults and late-instar nymphs) was used to assess the use of a wallow by red kangaroos. This is because adult ticks live for many years and so adults trapped in the field range in age from newly moulted nymphs to mature adults at least several years old. Adults and 4NN–5NN complete feeding within an hour or so of attaching to the host and so fall back into the original wallow, whereas larvae and 1NN and some 2NN feed for some days (Doube, 1972) and so are distributed sequentially in the series of kangaroo wallows occupied over the days succeeding attachment.

### 2. 4. Sampling tick populations in the field

Field populations of the tick were sampled in the study area at intervals over 3-years (February 1969 to March 1971). On each occasion, paired CO_2_ traps were set in kangaroo wallows under shade trees.

### 2. 5. Incidence of diapause in field-collected females

Female ticks were collected on 21 occasions between March 1968 and March 1971 and provided with an opportunity to feed on rabbits. Most diapausing females fed but did not produce eggs within the usual time frame (10–20 days at 30ºC) whereas some ticks, in deep diapause, refused to feed. Egg production was resumed in most females after a period of diapause development (Doube, 1972) [Each year most females (regardless of age) enter a non-reproductive diapause in which they feed but do not lay eggs (and eventually refused to feed) but then return to breeding condition for a period, during which time they have the capacity to lay several batches of eggs (Doube, 1972).]

### 2.6. The use of shade trees by red kangaroos

Kangaroo wallows under trees in the main study area were examined about every two months over an 18-month period (October 1969 to March 1971). The frequency of use of wallows by kangaroos was estimated by recording the amount and type of dung present in the wallows under each tree. Dung was classed by its colour and texture into two groups: old and new. If both old and new dung classes were present the wallow was recorded as having been visited at least twice. After each wallow was examined, all the kangaroo dung was removed. Because the intervals between wallow examinations were not equal, the number of wallows visited by kangaroos was not used as a comparative statistic and instead a ‘wallow-use index’ was devised to take account of the unequal time intervals between dung sampling. The wallow-use index = number of times wallows were visited x 10^4^ / total number of wallows x interval in days. For example, between 7 January and 18 March 1970 (an interval of 70 days), 169 wallows were used 122 times, and so the wallow-use index was 122 x 10^4^ / 169 x 70 = 104.

The quality of shade provided by each tree was allotted to one of three classes defined in terms of the intensity of light transmitted through the foliage on a bright, clear day when the intensity of direct sunlight was about 100,000 lux. The average of three readings under each tree was taken as the index of light intensity in the wallow. Low light intensity indicated dense shade. The three classes were: (i) dense shade – almost no direct sunlight reached the ground through the foliage (< 1300 lux); (ii) medium shade – some sunlight came directly through the foliage (1300 < Intensity ≤ 33,000 lux); and (iii) sparse shade – the foliage obscured very little of the sunlight (> 33,000 lux).

### 2.7. A secondary study

A secondary study was conducted in an adjacent valley on Moralana Station on a cloudless day in December 1970 with a maximum shade air temperature of 32°C. The shade trees in an area 1.8 x 0.32 km (1.1 x 0.2 miles) were mapped and classified according to the tree species and the quality (amount) of shade provided and the number of kangaroo wallows under each tree. The area examined was adjacent to the main study area but had never been sampled for ticks before and so tick populations were unaffected by tick removal during previous samplings. The kangaroo wallows under 10 trees of each of five types were examined for ticks using two CO_2_ traps per tree. Three readings of light intensity were taken at each tree. At the same time, the area of shade that each tree provided was estimated (length x breadth). Trapping took place between about midday and 4.00 p.m.

### 2. 8. Longevity of ticks

Batches of 200 recently engorged larvae, 1NN and 4NN, were each confined in permeable mesh bags containing 500 g of dry wallow sand and buried 4–5 cm deep in a wallow in March 1970. Similar bags of sand+ticks were placed in an adjacent sand dune. Bags were retrieved on four occasions during the following 12 months and the live and dead individuals in each bag recorded before replacing the live individuals in the wallow/sand dune. [Previous laboratory analyses of the susceptibility of newly moulted nymphs to dry conditions demonstrated that they were extremely resistant to desiccation (surviving for >800 days at 30°C and 10% relative humidity with only two 7-day periods for rehydration in a saturated atmosphere) (Doube, 1972); rehydration of desiccated nymphs and adults extended longevity (Doube, 1972). Ticks burrow into soft friable sand (from kangaroo wallows) and reach their desired depth within 2 hours (Doube, 1972). Males and females reached mean depths ±SD) of 18 mm and 35 mm, respectively. Extensive laboratory analysis of the survival and longevity of eggs and newly hatched larvae under a range of temperature and humidity conditions showed that eggs and larvae were relatively vulnerable to arid conditions. Eggs hatched in 5.9 days at 35°C, which increased to 30.7 days at 20°C (Doube, 1972). The mean longevity of solitary larvae varied from 3.0 days at 35°C and 10% relative humidity to 127.5 days at 20°C and 95% relative humidity (Doube, 1972). Periodic rehydration did not increase longevity. However, solitary eggs are not evident in the field, where the egg batches remained as an egg mass under the female. Further, the hatched larvae remain clumped under the female until disturbed by CO_2_, when they emerge to seek a host. Fed females were allowed to oviposit buried beneath 2 cm of dry sand at 10% relative humidity and 30°C and 35°C. Larval survival was checked after 15 and 20 days.]

## 3. Results

### 3. 1. Age structure of tick populations in kangaroo wallows

In an initial study using paired kangaroo wallows under each of six separate shade trees, two traps were set in the morning, and the ticks collected six hours later (Table 1). The number of ticks caught in adjacent traps within wallows under the same tree were very similar to each other, indicating that the ticks in each trap were representative of the population available to be sampled in wallows under the same tree. There was a moderate degree of variability between wallows in the mean number of large ticks (adults + 3–5NN: range 15 to 70 per trap per wallow) but a very wide degree of variability in the numbers of small ticks (1NN+2NN) (range 3 to 359 per trap per wallow). The variance ratio of the two samples (F=15.6) indicated a highly significant difference in the level of variability between the two samples.

**Table 1.** The incidence of occurrence of small (1NN and 2NN) and larger ticks (3–5NN and adults) in paired traps in wallows under six shade trees. 1NN, first-instar nymphs; 2NN, second-instar nymphs; 3–5NN, third to fifth-instar nymphs.

|  | 1NN and 2NN |  | Adults & late-instar nymphs |  |
| --- | --- | --- | --- | --- |
|  | Trap 1 | Trap 2 | Trap 1 | Trap 2 |
| Tree 1 | 6 | 3 | 14 | 15 |
| Tree 2 | 10 | 12 | 59 | 58 |
| Tree 3 | 34 | 41 | 18 | 12 |
| Tree 4 | 4 | 2 | 12 | 18 |
| Tree 5 | 42 | 60 | 27 | 20 |
| Tree 6 | 230 | 288 | 34 | 107 |
| Summary | Mean | 61.0 |  | 32.8 |
|  | SD | 95.2 |  | 28.6 |

### 3. 2. The use of wallows by kangaroos

The use of wallows by kangaroos varied throughout the year, as indicated by the quantity and quality of the dung present, with many wallows being used in the hot summer months (mean maximum temperature at nearby Hawker, 32–35^°^C) whereas few wallows were used in the cooler months (mean maximum of 10–13^°^C) (Fig. 3).

**Fig. 3.**
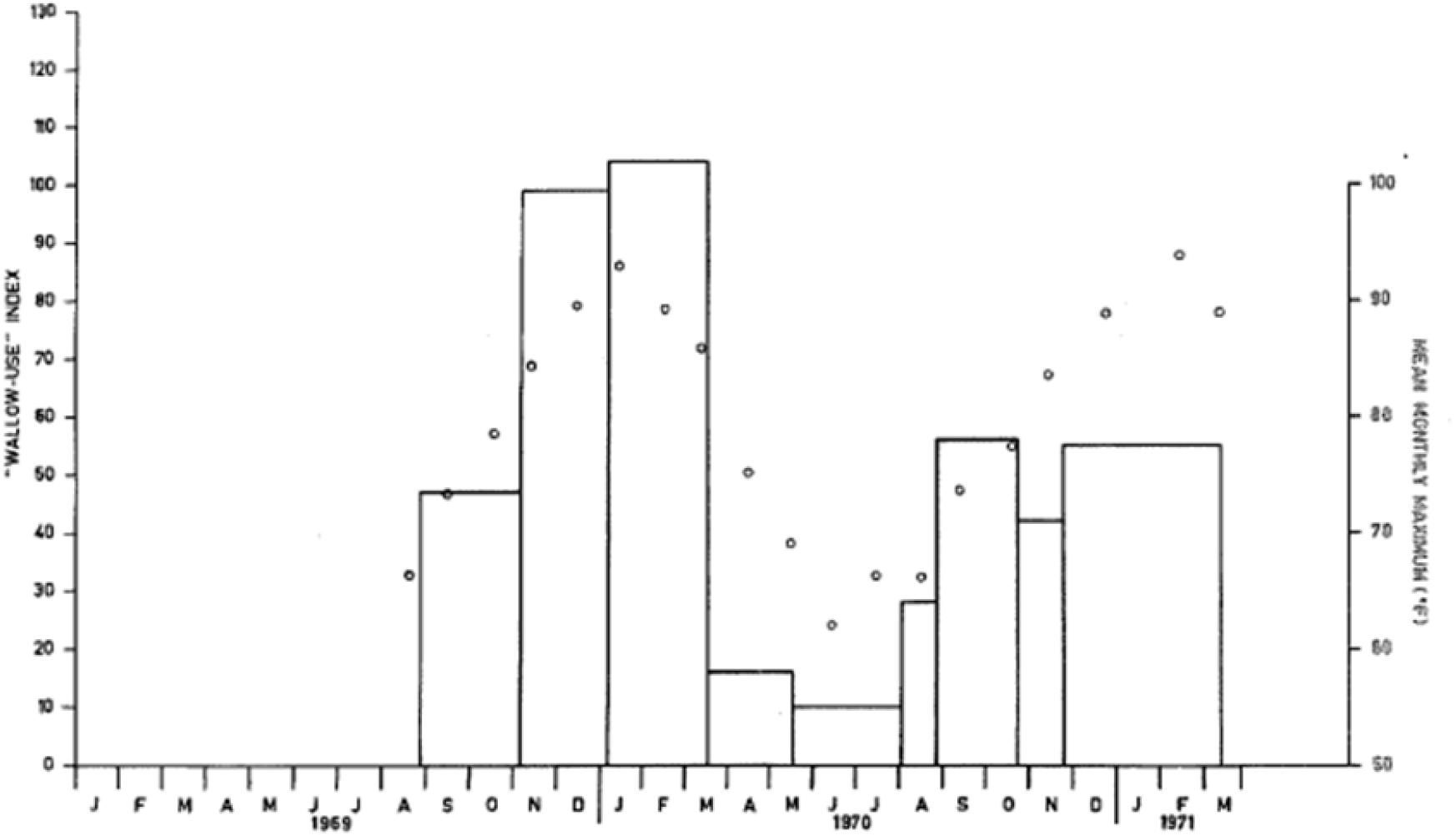
Kangaroo wallow-use index, seasonal changes in the frequency with which wallows were used by kangaroos; versus mean maximum monthly temperature at nearby Hawker, South Australia. The log of the number of ticks found was plotted against a shade-use index (the product of the three light intensity readings).

A supplementary experiment examined the relationship between age structure of the ticks or tick numbers, and the density of the shade and the area of wallows on one day in December 1970. There was no effect of the density of the shade, or the area of the wallows, on the age structure of the tick population, with about 10–20% as 1NN, 30–40% as 2NN, 20–30% as 3NN–5NN and 20–30% as adults (Table 2). In contrast, there was a major effect of shade density on tick numbers, with the trees providing the densest shade harbouring the most ticks (Fig. 3). For example, 10 times more ticks were trapped in wallows under dense shade than in wallows under sparse shade (37.8 vs 4.0 ticks per wallow, n=14 and 12 wallows, respectively). There was a significant effect of wallow size on the number of ticks trapped, with the smaller wallows (average size 0.6 sq m) harboured substantially fewer ticks (mean 6.0 ticks per wallow) than did the larger wallows (Table 2).

**Table 2.**
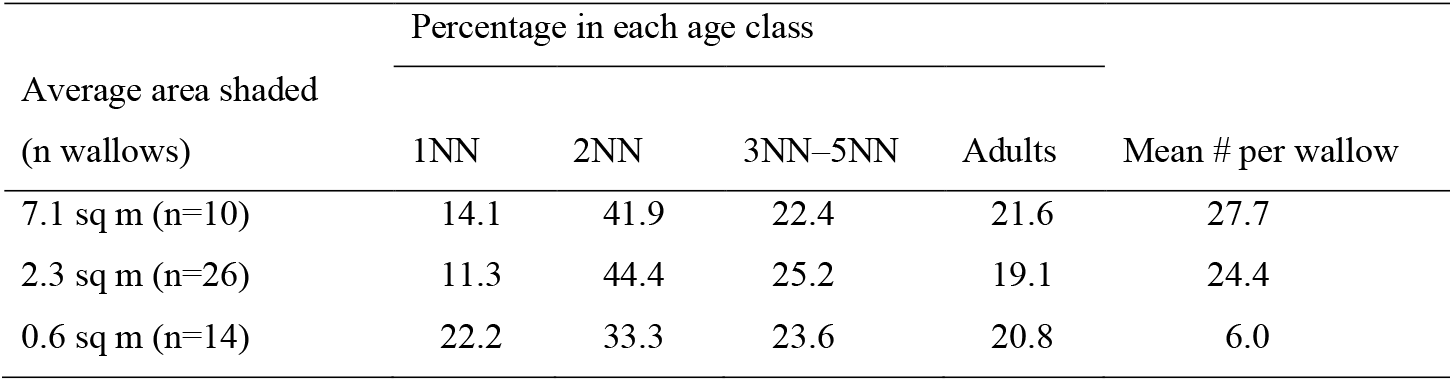
The effect of area of shade provided by trees on the age structure and mean size of populations of the tick *Australpavlovskyella gurneyi*. In December 1970, paired traps were set under 10 of each of the five dominant tree species in the study area. The traps were baited in the morning and cleared in the afternoon.

These data suggest that kangaroos begin to wallow frequently when the mean monthly maximum is about 21–27°C and prefer larger trees that provide ample dense shade.

Resting in the shade serves a thermoregulatory function for kangaroos so it is likely that the quality of the shade chosen varies with temperature and hence with the season of the year. To examine this possibility the data in Fig. 3 were reanalysed and the number of kangaroo wallows of a particular shade class that had been used was expressed as a percentage of the total number of wallows that had been used. This is an index of the class of shade chosen by the kangaroos.

The proportion of kangaroo wallows in each shade class used during the cooler months (May to August) was about equivalent to the proportion of trees in that class (Table 3). Thus, during the cooler months of the year the kangaroos showed no preference for any particular shade quality.

**Table 3.**
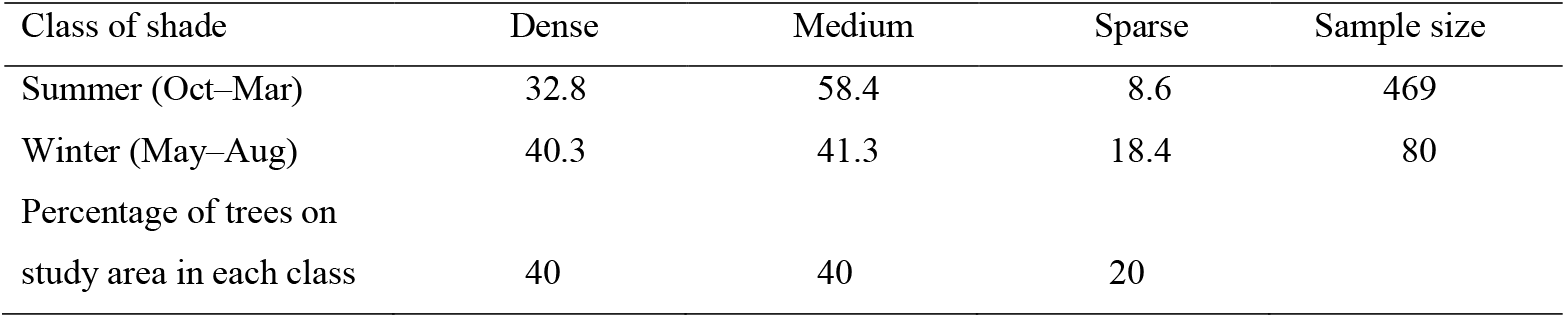
Percent use of shade by kangaroos for three classes of shade (dense, medium or sparse), calculated by recording the number of times wallows under each class of shade were visited by red kangaroos. This was expressed as a percentage of the total number of visits to wallows by the kangaroos.

| Class of shade | Dense | Medium | Sparse | Sample size |
| --- | --- | --- | --- | --- |
| Summer (Oct–Mar) | 32.8 | 58.4 | 8.6 | 469 |
| Winter (May–Aug) | 40.3 | 41.3 | 18.4 | 80 |
| Percentage of trees on<br>study area in each class | 40 | 40 | 20 |  |

### 3. 3. Larval and nymphal longevity

The strategy of clumping of eggs and larvae under a female until disturbed (also known as female brooding behaviour) increases the longevity of larvae (Table 4). Mean longevity (± SD) of solitary larvae at 10% RH and 30^°^C and 35^°^C was 3.9 ± 1.0 and 3.0 ± 0.6 days (Doube, 1972). Individual batches of eggs under females gave rise to clumps of eggs/larvae (mean ± SD = 308 ± 180 eggs per female). Egg hatch and survival for eggs/larvae clumped under females was assessed after 15 and 20 days at 10% RH and 30°C and 35°C. After 20 days at 30°C, most of the eggs had hatched and nearly all larvae were still alive. In contrast, hatching and survival at 35°C was seriously reduced but still greater than that observed with solitary larvae at 35^°^C. Clearly the clumping of larvae under the females seriously increased the protection of larvae from desiccation.

**Table 4.** The survival of eggs and larvae of *Australpavlovskyella gurneyi* clumped under female ticks buried in 2 cm of dry sand and held at 10% relative humidity and at 30° C and 35°C.

| Exposure temperature | Mean % hatch (±SD) |  | Mean % alive (±SD) |  |
| --- | --- | --- | --- | --- |
|  | 30°C | 35°C | 30°C | 35°C |
| 15 days | 83.4(±7.7) | 26.0 (±4.2) | 99.5 (±1.0) | 60.5 (±2.3) |
| 20 days | 79.0(±1.0) | 7.3 (±4.5) | 96.7 (±1.0) | 2.3 (±4.0) |

Longevity in the absence of a meal was tested in the field at Moralana Station where engorged larvae, 1NN and 4NN were placed in permeable chambers (gauze bags) in autumn in a wallow and in an adjacent sand dune. All moulted into the next instar and mortality was assessed at intervals over the following 52 weeks (Table 5). Most 2NN and adults were able to survive for at least 12 months in the absence of a meal.

**Table 5.** Effects of exposure to field conditions on the survival of engorged larvae, 1NN and 4NN of *Australpavlovskyella gurneyi* placed at a depth of 5–8 cm in sand in a kangaroo wallow and an adjacent sand dune on Moralana Station in March 1970 and assessed at intervals during the following year. There were 200 individual ticks per treatment.

| Exposure (weeks) | Percent mortality |  |  |
| --- | --- | --- | --- |
|  | Larvae | 1NN | 4NN |
| 9 in wallow | 9.5 | 0.5 | 0 |
| 20 in wallow | 13.0 | 0.5 | 0 |
| 32 in wallow | 22.0 | 6.0 | 1.0 |
| 42 in wallow | 31.5 | 7.0 | 1.0 |
| 52 in wallow | 41.0 | 15.5 | 6.0 |
| 52 in sand dune | 100 | 98 | 27 |

### 3.4. Larval production and adult diapause

Larvae were present in large numbers during early summer but scarce or absent during autumn and winter (Table 6). The incidence of reproductive diapause in field-collected females was assessed in the laboratory on 21 occasions from April 1969 to March 1971. The data (pooled on a one-year x-axis: Fig. 4) indicates a strong seasonal cycle in the incidence of diapause in which 80–100% of the individuals collected between January and May failed to produced eggs (i.e. were in diapause), whereas during October to mid-December at least 95% of all females produced eggs and so were not in diapause (Fig. 4).

**Table 6.** Changing annual incidence of larval *Australpavlovskyella gurneyi* in wallows of red kangaroos on Moralana Station.

| Data of sampling | # wallows with larvae (sample size) | Mean # larvae per positive sample |
| --- | --- | --- |
| 20 March 1971 | 6 (10) | 72 |
| 20 December 1970 | 12 (12) | 40 |
| 28 August 1970 | 0 (12) | 0 |
| 1 August 1970 | 0 (12) | 0 |
| 6 May 1970 | 9 (10) | 3 |
| 17 March 1970 | 5 (8) | many |
| 7 January 1970 | 5 (8) | many |
| 8 November 1969 | 1 (4) | 18 |
| 24 August 1969 | 8 (9) | 1 |
| 4 May 1969 | 6 (7) | 0 |
| 4 April 1969 | 3 (5) | 0 |

**Fig. 4.**
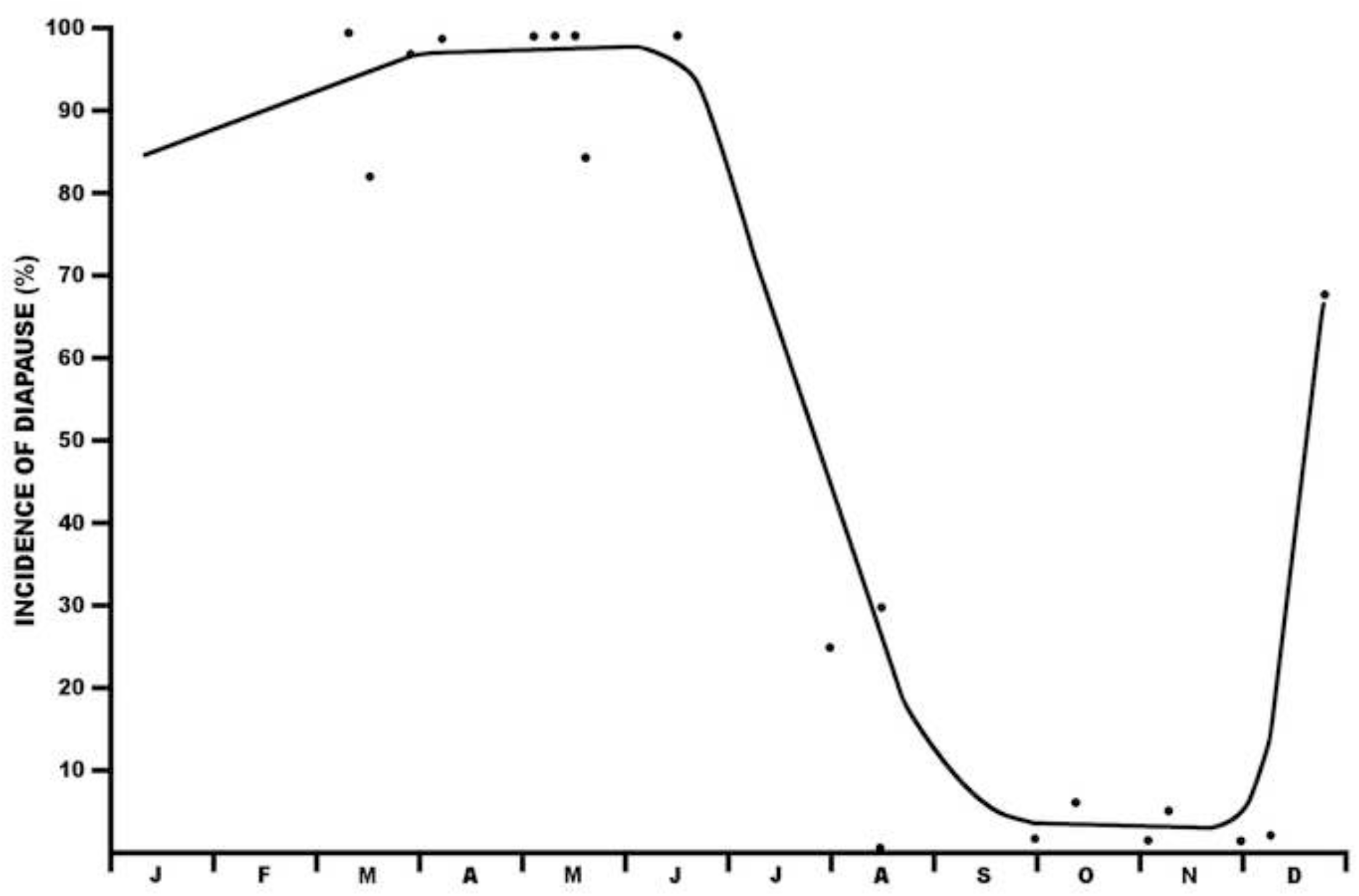
The incidence of adult reproductive diapause in *Australpavlovskyella gurneyi* collected from the field between January 1969 and March 1972.

### 3.3. Population processes over 3 years

The percentage of the tick population represented by first-instar nymphs ranged from about 10% to 40% depending upon the season of the year, with the higher percentages in late summer, following a spring–summer incidence of larvae. This was followed in autumn by an increase in the incidence of 2NN in early autumn (Fig. 5).

**Fig. 5.**
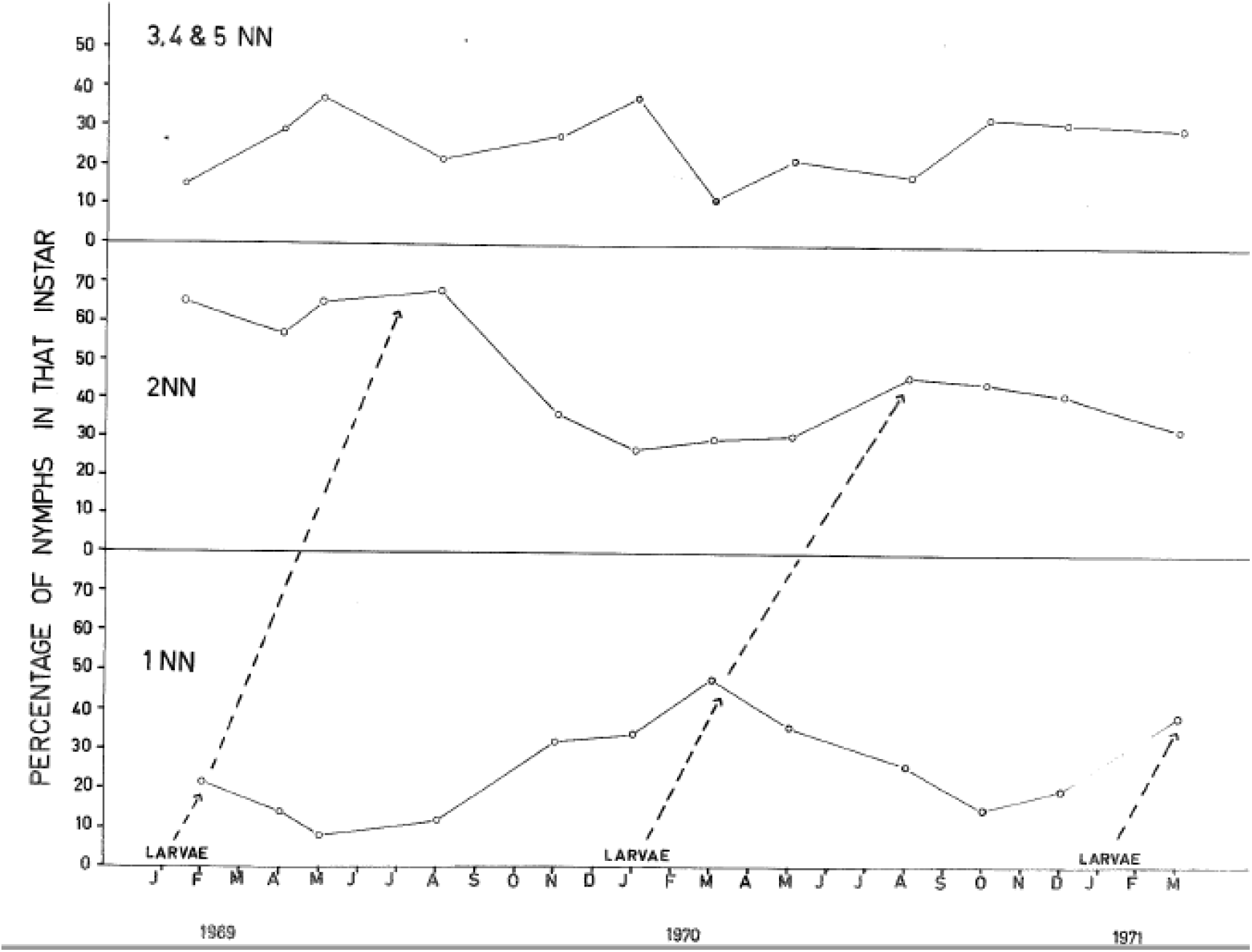
Fluctuations in the relative proportions of first- and second-instar-nymphal *Australpavlovskyella gurneyi* and the third to the fifth-instar nymphs over a 26-month period, from 111 kangaroo wallows and > 6000 ticks.

## 4. Discussion

The overriding feature of the environment of the kangaroo soft tick, *A. gurneyi*, is its harshness and unpredictability, but this threat is reduced by the alignment of the biology of the tick with the ‘wallowing’ behaviour of the red kangaroo. The red kangaroo shows no shade preference during winter but preferentially chooses isolated large shade trees providing medium or dense shade under which to shelter from the heat of the day in summer (Fig. 3, Table 2) (Browning, 1962). This activity produces sandy depressions, or wallows, under appropriate shade trees, in which the tick lives when not feeding. Although most kangaroo wallows on Moralana Station were visited by kangaroos only 2–3 times per year, this provided ample food to support a large tick population.

Dispersal between kangaroo wallows is achieved by way of a circadian rhythm of detachment of engorged larvae, 1NN and 2NN which ensures that engorged ticks detach in the middle of the day (when the kangaroo is resting in the shade) some days after first attaching (Doube, 1975d).

The instar distribution within any wallow is very variable due to the feeding behaviour of the tick and the behaviour of’ its host. For example, on one day a red kangaroo might rest in a wallow and in so doing remove a proportion of the larvae and 1NN from the wallow. These larvae and nymphs will then be deposited in relatively large numbers in a small number of kangaroo wallows over the following few days. Thus, the instar distribution of several wallows might be dramatically altered in a few days and then perhaps remain unaltered for months. This seems to be illustrated by the data in Table 1. The coefficient of variation for 1NN+2NN was far greater than that for 3NN–5NN and adults.

In order to appreciate the host : parasite interaction, one must also understand the patterns of movement and the daily routine of the host, the red kangaroo. Frith and Calaby (1969, p. 87, p.96) said:

Red kangaroos are nomadic and their distribution, at any time, is determined by the availability of green food and perhaps, to some extent, by shade also. The scale of movement of individuals is related to the severity of drought and the distance that must be travelled to find green food … when these (food and shade) are abundant only local movements [several miles] are necessary and larger ones are not undertaken. In times of stress, however, the animals have the ability, and the stamina to move considerable distances in search of it (sic).

Further, when discussing the daily routine of the kangaroo, Frith and Calaby (1969, p. 61) said:

In hot periods they spend the days under shady trees and move out to feed and water at dusk, returning at dawn … During the day, the animals camp in dust baths dug under bushes or shrubs and what seem to be the same animals return to the same spot day after day while the mob remains in that general area.

Our findings, based on observations in a particularly arid environment, were somewhat at variance with this view in that we rarely found large quantities of dung in individual kangaroo wallows and commonly there was evidence of only one visit to a wallow during the sampling intervals, commonly of 1–2 months. This statistic provides the basis of our interpretation of the ecology of the red kangaroo tick in severe desert environments.

### 4.1. Habitat and seasonal dynamics of tick populations

During hot weather, red kangaroos prefer wallows under trees that provide large areas of dense shade. One consequence of this is that the density of ticks trapped under trees that provide the densest shade was substantially greater than under trees where shade was sparse (38.6 and 4.0 ticks respectively per wallow in January 1970; 2 traps per tree). Further, an average of 27.7 ticks per tree were trapped under the 10 trees with the largest shade area (an average of 76 sq. ft (7 sq. m) of shade), whereas an average of 6.0 ticks per tree were trapped under the trees with the smallest area of shade (an average of 6.3 sq. ft (0.6 sq. m) of shade). Thus, there were many more ticks present under trees that provided large areas of dense shade than under small trees that provided sparse shade.

The seasonal dynamics of the tick populations at Moralana Station are clearly dictated by the wallowing activities of the resident kangaroos. In summer, kangaroo wallows providing the best shade and the largest area shaded are preferred above others but during winter the kangaroos show no preference for shade intensity (Table 2). The high incidence of larvae in November–January (Table 6) gave rise to a peak in the proportion of 1NN somewhere between January and March and this flush of 1NN gave rise to a slight rise in the proportion of 2NN somewhere between March and April (Fig. 3). There appears to be no seasonal trend in the proportion of 3–5NN or adults in the population.

Although favoured kangaroo wallows on Moralana Station may be visited as often as three or four times during the hot period of the year, as previously indicated, each tick in the wallow is unlikely to feed that many times. Since the tick requires 5–7 meals to complete its life cycle, it is unlikely that the average life cycle will be completed in under 2–3 years, even under favourable conditions, and it may take 5–10 years under unfavourable conditions.

### 4.2. Reproduction–diapause

During spring and early summer, most female ticks have a short pre-oviposition period (<20 days at 30°C) but in late summer the preoviposition period becomes more variable and sometimes extends to 50 days. This results in a flush of short-lived larvae in early summer (when the kangaroo wallows are being visited relatively frequently by kangaroos). Late-summer and autumn larvae come from a small proportion (10–15%) of females that do not enter diapause or have an extended pre-oviposition period, resulting in a trickle of larvae appearing until diapause intervenes to inhibit oviposition in late autumn. Many females engorge in one year but lay eggs in the spring of the following year after diapause development has been completed, obviating the need for an initial meal in spring.

### 4.3. What is the chance of *Australpavlovskyella gurneyi* finding a meal before it dies?

A number of factors affect the probability that a tick will find a meal in any one year. First, a kangaroo must visit the wallow in which the tick is living. Second, the tick must be capable of sensing and responding to the sensory cues that the presence of a kangaroo provides. Third, the ticks must survive the interval between kangaroo visits, and fourth, the ambient temperature must be suitable for tick activity.

#### 4.3.1. Seasonal activity of kangaroos

On Moralana Station, kangaroos appear to visit shade trees to shelter from the heat primarily during the warmer seasons of the year (Fig. 3) and wallows appear to be visited, on average, only 2–3 times during that period.

#### 4.3.2. Population responses to CO_2_

Neville (personal communication) found that the response to CO_2_ of field populations of *O. savignyi* varied widely between individuals. Some responded immediately whereas others failed to respond over a 2-year period, during which the sand in which they lived was regularly sampled using CO_2_ as a bait. *A. gurneyi* responded in a similar fashion to traps baited with CO_2_ set in a wallow (fenced to exclude kangaroos) on each of 10 successive sampling trips over a period of two years. Ticks were trapped on the first eight occasions. There was no evidence (neither dung nor scratchings) that the wallow was visited during that period and, altogether, about 800 ticks were taken from the wallow. It seems therefore likely that only a relatively small proportion of the ticks in a wallow feed each time a kangaroo visits it. This contrasts with the findings of Dupraz et al. (2017), who found that populations of *Alectorobius maritimus* were seriously depleted by successive trapping events.

#### 4.3.3. Longevity of larvae, nymphs and adults in the field

Solitary newly hatched larvae were relatively vulnerable to arid conditions in the laboratory, with the mean longevity of larvae varying from 3 to 128 days. Periodic rehydration did not increase longevity (Doube, 1972). However, solitary eggs are not present in the field, where egg batches and massed hatched larvae remain *en masse* under the female, thereby substantially increasing larval longevity. Nevertheless, larval longevity is too short to survive the cooler months of the year during which kangaroo wallows are rarely visited by kangaroos.

In marked contrast, about 60% of 1NN exposed to ambient conditions in a wallow on Moralana Station were still alive after 12 months without food, and most of the 2NN and 3NN survived similar exposure (Table 2). In the laboratory 2NN and 4NN survived for a long period (at least 800 days and 1140 days, respectively) in dry conditions (30°C and 10% relative humidity) with only two periods of rehydration and no feeding (Doube, 1972). Clearly, the larger ticks have a remarkable capacity to survive long periods without feeding and thereby persist during periods when red kangaroos are absent from a region for a year or two.

The capacity of nymphs and adults to withstand drought is enhanced their capacity to absorb water from unsaturated air. Rainfall records for nearby Yudnapinna Station (–32.06904, 137.05427) for the years 1855 to 1955 (Bureau of Meteorology and records from the Waite Agricultural Research Institute) indicate that an average annual rainfall of 200 mm (an annual average of 57 rainy days) was dispersed among all months of the year, with a predominance of light and very light showers interwoven with significant and heavy falls. Laboratory studies (Doube, 1972) indicate that moderate rainfall (3.5 mm) was sufficient to moisten the soil profile and allow ticks to rehydrate and so it seems likely that desiccation may be of minor importance in regulating tick populations in areas visited intermittently by kangaroos.

#### 4.3.4. Temperature dependent responses

The physiological condition of the tick must be such that it is capable of responding to the presence of a kangaroo (see above) and the ambient temperature must be greater than the temperature threshold for activity. The fact that all nymphal instars and adults were trapped at all times of the year indicates that, on Moralana Station, ambient temperatures during winter did not inhibit activity. Because the interval between kangaroo visits is often quite protracted, ticks have almost invariably moulted and are ready to feed again long before the kangaroo wallows are visited again, and so the effect of ambient temperature on the rate of development rarely, if ever, limits the rate of population growth.

### 4.7. Minimising the risk of local extinction

In the years leading up to 1959, red kangaroos became increasingly abundant throughout inland Australia, so much so that they were considered by some to be in plague proportions in the 1950s (Frith and Calaby, 1969). However, during the period 1959–66 the numbers of red kangaroos in many areas were greatly decreased by shooting for meat and by drought (Frith and Calaby, 1969). The effect of this reduction in kangaroo numbers on the tick population was evident in 1968 when the present study began. In the early 1960s the ticks were abundant and readily caught in many places in inland South Australia (Browning, 1962; Browning, personal communication) but by 1968 ticks were virtually absent from the same areas, even though occasional red kangaroos were seen in those areas (Doube, unpublished data). Although the tick population had crashed dramatically, there remained a few isolated pockets in which kangaroos and ticks were abundant. For example, on Moralana Station (due to the management’s ban on kangaroo shooting) kangaroos were still abundant and so the tick was still present in relatively large numbers [Kangaroos were still abundant on Moralana Station on the 6^th^ July 2019 when the two authors visited to the study site; ticks were also found at that time].

*Australpavlovskyella gurneyi* is an opportunistic animal adapted to surviving in the Australian outback, a harsh and unpredictable environment. This is achieved by spreading the risk of local extinction in time and space over a range of phenotypes, with variations in the proportion of ticks that respond to the presence of a host, in the proportion entering diapause and the depth of the diapause, and in their capacity to survive desiccation (Doube, 1972). In any season only some the mechanisms will have survival value, but over a long period each mechanism contributes to the maintenance and stability of the population, thereby reducing the probability of local extinction.

## Acknowledgements

The supervision provided by Professor Tom Browning and Dr Roger Laughlin of B.M. Doube’s PhD, is gratefully acknowledged. Loene Doube maintained the laboratory colonies of *A. gurneyi* and its hosts during my regular absences and provided valuable feedback on the structure and content of this manuscript. Dr Kerrie Davies provided substantial assistance with preparing the original figures. Ernest Teo provided assistance in updating the figures. The project was funded by a University of Adelaide PhD Research Grant (URG). We thank two anonymous referees.

